# Evidence for an aquatic origin of influenza virus and the order *Articulavirales*

**DOI:** 10.1101/2023.02.15.528772

**Authors:** Mary E. Petrone, Rhys Parry, Jonathon C. O. Mifsud, Kate Van Brussel, Ian Vorhees, Zoe T. Richards, Edward C. Holmes

## Abstract

The emergence of novel disease-causing viruses in mammals is part of the long evolutionary history of viruses. Tracing these evolutionary histories contextualises virus spill over events and may help to elucidate how and why they occur. We used a combination of total RNA sequencing and transcriptome data mining to extend the diversity and evolutionary history of the order *Articulavirales*, which includes the influenza viruses. From this, we identified the first instance of *Articulavirales* in the Cnidaria (including corals), constituting a novel and divergent family that we tentatively named the *Cnidenomoviridae*. This may be the basal group within the *Articulavirales*. We also extended the known evolutionary history of the influenza virus lineage by identifying a highly divergent, sturgeon-associated influenza virus. This suggests that fish were among the first hosts of influenza viruses. Finally, we substantially expanded the known diversity of quaranjaviruses and proposed that this genus be reclassified as a family (the *Quaranjaviridae*). We find evidence that vertebrate infecting *Quaranjaviridae* may have initially evolved in crustaceans before spilling into terrestrial Chelicerata (i.e., ticks). Together, our findings indicate that the *Articulavirales* has evolved over at least 600 million years, first emerging in aquatic animals. Importantly, the evolution of this order was not shaped by strict virus-host codivergence, but rather by multiple aquatic-terrestrial transitions and substantial host jumps, some of which are still observable today.

## INTRODUCTION

Zoonotic viruses transmitted from other animals to humans have caused multiple epidemics in recent decades^1,2^, and the frequency of these events is projected to increase as a consequence of climate change^3^. To combat this threat, research on zoonotic risk predominantly aims to identify viruses that are primed to spill over into humans. It is commonly assumed that the viruses most likely to cause epidemics in the future are genetically similar to those that have caused outbreaks previously^4,5^. A number of field-based and bioinformatic methods aimed at detecting potentially zoonotic viruses are predicated on this assumption^6–8^. Although these approaches help construct a picture of virus diversity, the process of crossspecies virus transmission that drives disease emergence in the short-term also plays a key role in virus speciation in the long-term, shaping virus-host associations that likely date back to the initial existence of single-celled organisms^9^. In addition to revealing their antiquity, elucidating the deep evolutionary history of known disease-causing viruses provides important context for understanding the true rate at which viruses jump species boundaries to infect new hosts.

Because they reveal the viromes of a diverse array of organisms, metagenomic data are a powerful tool for tracing long evolutionary histories^10^. These data have already shown that the global virosphere is vast and largely unexplored^10^, and a recent emphasis on the exploration of marine environments has demonstrated that the ocean is a rich source of virus diversity^11^. Metagenomic research has also shown that viruses once thought to be restricted to mammals exist in a wide variety of other vertebrates^12^, suggesting that they have a long evolutionary history in animals. Similarly, disease-causing vertebrate viruses have origins in invertebrate hosts. For example, members of the family *Flaviviridae* (single-strand positive-sense RNA viruses) were recently discovered in basal invertebrates (Cnidaria)^13^, basal chordates (Ascidia)^13^, and in diverse aquatic environments^14^. This suggests that the *Flaviviridae*, which include such human pathogens as dengue virus and yellow fever virus, emerged concurrently with the origin of the Metazoa (i.e., animals) some 750-800 million years ago^15^. Indeed, that aquatic ecosystems may have influenced the evolution of terrestrial viruses speaks both to the virus diversity these environments support and to their role as the source of life on Earth.

The order *Articulavirales* is of special importance in exploring the intersections of marine life and the evolution of disease-causing viruses. Viruses within the *Articulavirales* have segmented, negative-sense RNA genomes. This order is currently organised into two families – the *Orthomyxoviridae* and the *Amnoonviridae*. The former receives frequent public health attention because it contains three genera that can cause disease in humans: *Thogotovirus, Quaranjavirus*, and *Influenzavirus*. Thogotoviruses were first isolated in the 1960s^16^, while discoveries of disease-causing quaranjaviruses are ongoing (e.g., the discovery of Wellfleet Bay virus as the cause of avian mortality in 2014^17^). Both thogotoviruses and quaranjaviruses periodically spill over into humans, other mammals, and birds through tick-mediated transmission. In contrast, influenza viruses cause seasonal epidemics in human populations, as well as occasional pandemics.

Aquatic animals serve as hosts for both families. The *Amnoonviridae* is a family of primarily fish-infecting viruses^18^, and two species have been found in amphibians^19,20^. Tilapia lake virus, a genus within this family, has important implications for the global aquaculture industry because it is associated with severe disease in tilapia^21^. Similarly, salmon isavirus (*Orthomyxoviridae*) causes overt disease in salmon^22^, and influenza-like viruses have been identified in fish and amphibians^12,23^, although with unknown disease associations. Given their known host distribution, the *Amnoonviridae* likely emerged in aquatic animals, but the long branch lengths associated with this family in current *Articulavirales* phylogeny suggests substantial unsampled diversity.

Genomic architecture varies within and between the families of the *Articulavirales*, but all contain three polymerase segments: polymerase basic 1 (PB1), polymerase basic 2 (PB2), and polymerase basic 3/polymerase acidic (PB3/PA). These proteins form a heterotrimer comprising the canonical RNA-dependent RNA polymerase (RdRp)^24^ that is routinely used as a phylogenetic marker for RNA viruses^25^. PB1 contains the palm domain, defined by the SDD amino acid motif, and is therefore the most highly conserved of the subunits. For this reason, PB1 is the most likely of all the segments to be identified from metagenomic data.

We applied a combination of total RNA sequencing and data mining to revisit the evolutionary history of the *Articulavirales*. No *Articulavirales* genera have been discovered in basal non-bilaterian (i.e., lacking body symmetry) invertebrates of the Cnidaria, which includes corals, jellyfish, and hydra. To address this, we analysed the RNA viromes of Hexacorals and Octocorals from Australian. Concurrently, we screened the NCBI Transcriptome Shotgun Assemblies (TSA) database for novel orthomyxo-like viruses associated with aquatic animals: Phylum Arthropoda (e.g., shrimp and crabs); Phylum Chordata, Subphylum Vertebrata (fish); Subphylum Tunicata (sea squirts and tunicates); and Phylum Mollusca (e.g., squid and whelk). We combined the resulting data sets to characterise the role of ancient aquatic animals in the evolution of the *Articulavirales* and determine when this important order may have emerged in relation to the evolution of the Metazoa.

## RESULTS

### Discovery of highly divergent *Articulavirales* in basal invertebrates

Influenza viruses are overrepresented in publicly available data for viruses in the order *Articulavirales*. Of the 76,887 *Articulavirales* PB1 segments available on NCBI Virus in January 2023, only 291 (0.38%) were not influenza viruses (**Fig. 1a**). Thus, our first objective was to expand the known diversity of non-influenza viruses in this order.

**Figure 1.**
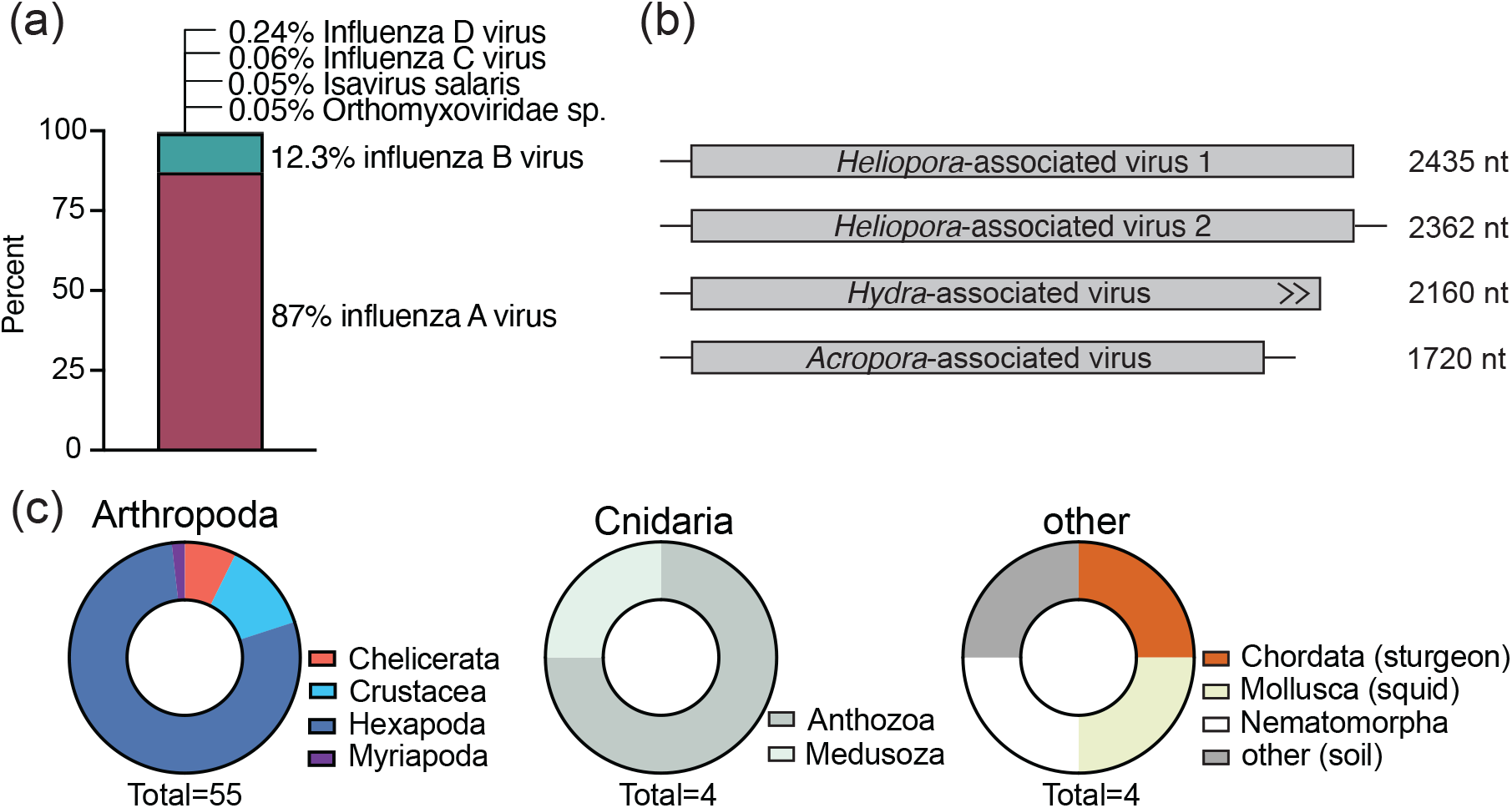
Expanding the known diversity of the *Articulavirales*. (a) Overrepresentation of Influenza A and B viruses in publicly available genomic data. Distribution of 99.7% of publicly available PB1 segments on NCBI Virus as of January 2023 by virus species. Virus species not shown each comprise <0.05% of available data. (b) PB1 segments of novel *Articulavirales* identified in Cnidaria hosts. *Heliopora*-associated virus 1 was identified through sequencing while *Heliopora*-associated virus 2 was identified through data mining. (c) Distribution of host species and classes associated with the *Articulavirales* detected through total RNA sequencing and datamining.

To search for highly divergent *Articulavirales*, we performed total RNA sequencing on 128 corals (Cnidaria) collected in Western Australia. From these we identified *Articulavirales* PB1 segments in two libraries - *Heliopora coerulea* (blue coral) and *Acropora samoensis* (stony coral) - using Diamond BLASTx. Although PB1 is the most conserved polymerase segment, both segments were highly divergent, sharing ≤ 25% amino acid identity with those of known viruses (**Table S1**). Neither contained premature stop codons nor returned BLAST hits to known coral genes, suggesting they are not endogenous virus elements (EVEs). However, they were present at very low abundance (both <0.001% non-rRNA reads). The segment recovered from the *Acropora*-associated library (1720nt) was shorter than the *Heliopora*-associated segment (2435nt) (**Fig. 1b**), and the two segments shared minimal sequence similarity (17.8% amino acid pairwise identity). This substantial genetic distance is consistent with the phylogenetic relationship of *Heliopora* and *Acropora* corals. The former are commonly known as blue corals belonging to the Subclass Octocorallia, Order Scleralcyonacea, which are estimated to have split from other Anthozoa about 500 million years ago^26^. *Acropora*, commonly known as staghorn corals, are from the Subclass Hexacorallia, Order Scleractinia, which emerged more recently, likely in the Permian around 260 million years ago^26^. Analysis of the composition of each library showed that the majority of eukaryotic genetic material (98%) in the *Acropora* library were associated with Cnidaria, while 73% of contigs assembled from the *Heliopora* library were Suessiales (dinoflagellate symbionts of corals). In this library, most contigs associated with Anthozoa (76%) were assigned to Scleractinia rather than *Helioporidae*, suggesting contamination or misassignment (**Fig. S1**).

We used these segments to screen Cnidaria assemblies available in the NCBI Transcriptome Shotgun Assemblies (TSA) database (n = 50). This screen yielded two additional novel PB1 segments: one in a *Heliopora coerulea* coral library and another in a *Hydra vulgaris* hydra library. The second *Heliopora*-associated PB1 segment (2362nt) was slightly shorter than the first (2435nt) and shared 47.6% amino acid pairwise identity. The open reading frame (ORF) of the hydra-associated segment was uninterrupted but incomplete (**Fig. 1b**). PB2 and PB3/PA segments could not be identified in either.

Having identified *Articulavirales* in basal invertebrates, we reasoned that this viral order likely contains unrealised diversity among other aquatic invertebrate hosts. To address this, we used the PB1 segment of the Wenling hagfish influenza virus to screen TSA libraries from Arthropoda, Tunicata, Mollusca, and fish. We selected this virus because of its aquatic host - a jawless fish - and its basal position in the *Orthomyxoviridae* in the current *Articulavirales* phylogeny. Using this approach, we detected 30 novel PB1 segments of at least 1000nt in length, 29 of which were associated with Arthropoda hosts and one associated with a Siberian sturgeon (*Actinopterygii*, ray-finned fish). To supplement this data set, we evaluated all RdRp contigs assembled from the Serratus project with an assigned “unknown” *Orthomyxoviridae* RdRp palmprint as per the palmDB database^27^. After removing known viruses, duplicated sequences, and fragments less than 800nt in length, we identified 29 additional novel viruses. In total, we identified 63 novel *Articulavirales* PB1 segments. The majority of these viruses were associated with arthropod hosts (n = 55), which likely reflects a sampling bias towards Arthropoda rather than the true host distribution of *Articulavirales* (**Fig. 1c, Supp. Data**).

Previous studies have presented evidence for the endogenization of *Orthomyxoviridae* polymerase segments^28^. We therefore assessed whether the contigs identified here could be EVEs. PB1 candidates that included premature stop codons were not included in our data set. Because the presence of multiple virus segments in a library would be strongly indicative of exogenous expression, we screened each TSA library in which we had found a PB1 segment for additional virus segments. With the exception of the coral- and hydra-associated libraries, we recovered three polymerase segments for all of the novel viruses we identified, and for the majority (n = 45), we successfully retrieved nucleoprotein and haemagglutinin segments (**Supp. Data**). Our inability to detect additional polymerase segments in the basal invertebrate libraries was likely due to very low sequence similarity of these segments to those of any known virus.

### The *Articulavirales* comprises at least four families and likely emerged in ancient, aquatic animals

Given the low sequence similarity of the PB1 segments we identified in the Cnidaria libraries, we hypothesised that these viruses would be phylogenetically distinct from other families in the order. To test this, we aligned the 63 sequences identified in this study with a representative sample of publicly available *Articulavirales* PB1 sequences using MUSCLE^29^ and inferred a maximum likelihood tree using IQTree^30^.

Notably, the inclusion of our newly identified viruses changed the phylogenetic structure of this order. Our cnidaria-associated viruses formed a putative new family currently comprising non-arthropod invertebrates. We have provisionally named this family the *Cnidenomoviridae*. Known hosts of this family are coral, hydra, tunicate (Urochordata), sea cucumber (Echinodermata), and mussel (Mollusca)^31^ (**Fig. 2a**, *Cnidenomoviridae*). Viruses from *Heliopora coerulea* corals were more closely related to each other than to the virus identified in the *Acropora* coral, tentatively suggesting class specificity (**Fig. S2**). This relationship also indicates that *Heliopora* is the host of the *Heliopora*-associated virus we identified through sequencing despite the library composition. Interestingly, viruses associated with Mollusca hosts do not belong exclusively to the *Cnidenomoviridae*. In particular, we identified an orthomyxo-like virus associated with a squid (*Sepioloidea lineolate*) that forms a clade with viruses associated with fish and whelk (Mollusca) (**Fig. 2a, *gold arrow***). The placement of this clade, which comprises exclusively aquatic hosts (salmon louse, marine flatworm, lizardfish, squid, whelk, crustacean, gobie), is poorly resolved relative to the *Amnoonviridae* (ufboot = 56). Interestingly, there is no clear separation of vertebrate and invertebrate hosts as is observed in the rest of the order (e.g., *Thogotovirus* and *Influenzavirus*), although true host associations have not been verified. Robust sampling may reveal that this clade constitutes its own family. The addition of the arthropod-associated viruses identified through our TSA and palmDB screens suggests that quaranja- and quaranja-like viruses should be reclassified as a new family – the *Quaranjaviridae* (**Fig. 2a**). All known viruses in this family are associated with arthropods as the primary host. The tree topology also supports the reclassification of salmon isavirus, which is currently classified as *Orthomyxoviridae*, as a member of the *Amnoonviridae* (**Fig. 2a**, ***green arrow***). Thus, we propose that *Articulavirales* comprises at least four families: *Orthomyxoviridae, Amnoonviridae, Quaranjaviridae*, and *Cnidenomoviridae*. We adopt this new nomenclature for the remainder of the manuscript.

**Figure 2.**
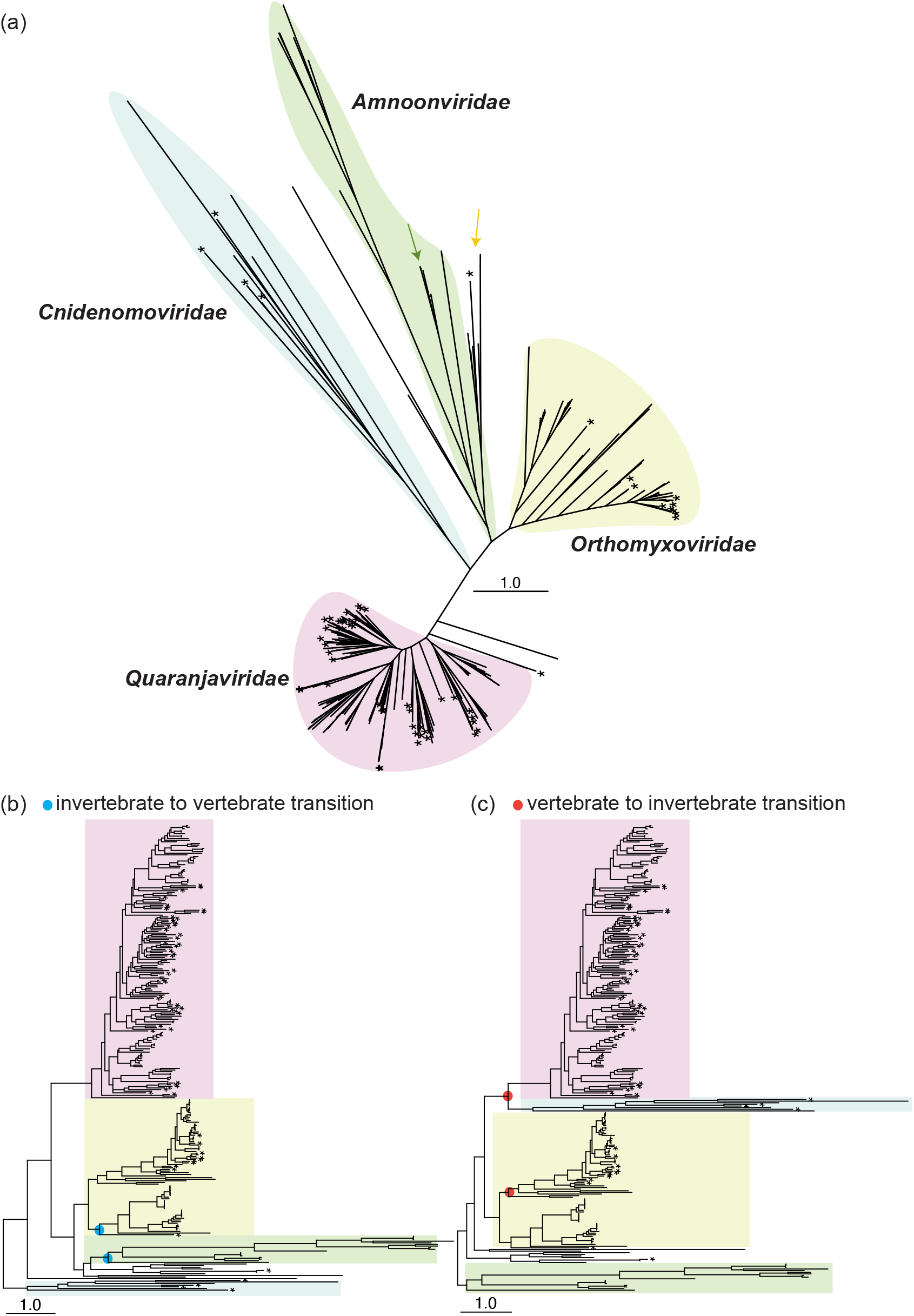
Phylogenetic evidence for four virus families in the order *Articulavirales*. (a) Unrooted maximum likelihood tree of the *Articulavirales* PB1 segment, with branch lengths scaled to the number of amino acid substitutions per site. *Novel viruses identified in this study. The green arrow marks Salmon isavirus, while the gold arrow marks an aquatic orthomyxo-like virus clade associated with Mollusca and fish hosts. (b) PB1 maximum likelihood tree rooted on basal invertebrate family *Cnidenomoviridae* (c) PB1 maximum likelihood tree rooted on the *Amnoonviridae*. In all cases branch lengths are scaled to the number of amino acid substitutions per site.

To interpret the evolutionary history of *Articulavirales*, we considered various possible rooting positions on the phylogeny. The structure of the tree presents ambiguities because both the *Amnoonviridae* and *Cnidenomoviridae* are defined by long branch lengths, and both are plausible roots. We first assessed whether the branching patterns were an artefact of an alignment error by comparing the topology of trees inferred using MUSCLE^29^ and MAFFT^32^ aligners, which apply different alignment algorithms (**Fig. S3**). No substantive differences were observable in tree topology, suggesting instead that large sampling gaps persist in these families and account for the branch lengths. Rooting the phylogeny on the novel basal invertebrate family *Cnidenomoviridae* is compatible with long-term virus-host codivergence and hence a reasonable hypothesis if the *Articulavirales* are indeed ancient (**Fig. 2b**). This phylogenetic history means that the *Articulavirales* first emerged in aquatic, sessile invertebrates, potentially around 600 million years ago when Medusoza and Anthozoa diverged^26^. The *Amnoonviridae* form a sister clade with *Orthomyxoviridae* and are basal to *Quaranjaviridae*, the former of which is the only other family known to include fish hosts. Under the parsimony criterion, this transition to vertebrate hosts would have occurred twice and independently (**Fig. 2b, *blue circles***). In contrast, placing *Amnoonviridae* as the root introduces two vertebrate-invertebrate transitions, contradicting virus-host co-divergence (**Fig. 2c**). In this representation of phylogenetic history, the *Cnidenomoviridae* are basal relatives of the *Quaranjaviridae*, and there is a clear division between invertebrate-only and vertebrate clades. There is also a division between arthropod-infecting (*Quaranjaviridae*) and nonarthropod-infecting (*Cnidenomoviridae*) viruses, although additional sampling may eventually refute these designations. In both cases, the tree structure indicates that *Articulavirales* first emerged in aquatic hosts before spreading to terrestrial animals.

### Fish were among the earliest hosts of influenza virus

Tracing the evolution of influenza virus could shed light on the mechanisms by which this virus emerged in mammals. In our TSA screen we identified the PB1 segment of a highly divergent influenza-like virus associated with the transcriptome of a Siberian sturgeon (*Acipenser baerii*) intestine (TSA ID: GIPE01). We successfully recovered the PB2 and PB3/PA segments from this library. All three segments contained complete ORFs (**Fig. 3a**). The genome of this virus differed from that of canonical influenza viruses, including those discovered in other fish, in two notable ways. First, the PB2 segment of this virus is approximately 100 amino acids longer than other segments in the clade (**Fig. 3a**). These additional amino acids do not appear to be the result of an insertion as they occur at 3’ the end of the segment. Second, while there is a conserved VGG motif in the palm domain of the PB1 segment of known influenza and thogotoviruses, the PB1 segment of the sturgeon-associated virus motif is IGG which is common among *Quaranjaviridae* and entirely absent in the *Amnoonviridae* (**Fig. 3a**). In contrast, polymerase motifs I-IV were well conserved (**Fig. 3a**). In Motif I, ‘KWNE’ is conserved throughout the clade. The hagfish and sturgeon influenza viruses share identical sequences in Motif II. While the SDD domain is requisite for functional *Articulavirales* PB1 segments, influenza viruses have a second serine (SSDD) in Motif III. Interestingly, Motif IV is more divergent in the hagfish virus compared to the sturgeon virus, but all are characterised by asparagine and serine in the third and fifth positions, respectively.

**Figure 3.**
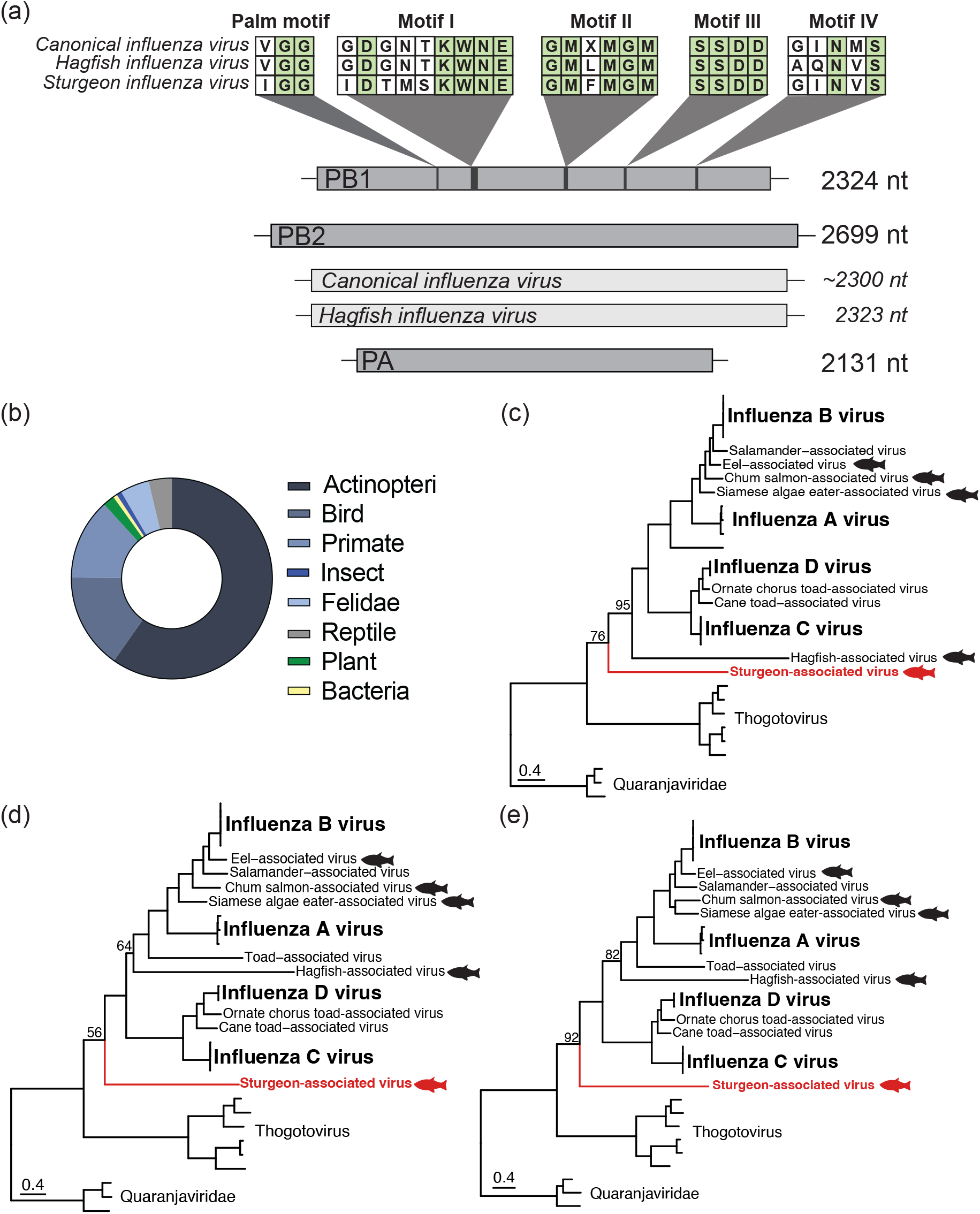
Characterisation of a divergent influenza-like virus identified in a Siberian sturgeon. (a) Structure and conserved motifs of the three polymerase motifs. (b) Composition of Siberian sturgeon intestine transcriptome evaluated using KMA v1.3.9a^33^ and CCMetagen v1.1.3^34^. Maximum likelihood phylogenetic trees of influenza PB1 (a), PB2 (b), and PB3/PA (c) segments. Fish hosts are indicated with fish icons. Segments from the novel sturgeon-associated virus are shown in red. All trees were rooted using the *Quaranjaviridae* as outgroups with branch lengths scaled to the number of amino acid substitutions per site.

As the sturgeon transcriptome comprised the animal’s intestine, we hypothesised that this virus was likely derived from the animal’s diet. To test this, we analysed the library composition using KMA v1.3.9a^33^ and CCMetagen v1.1.3^34^. Interestingly, 64% of all contigs were associated from the host (Actinopteri), and 29.5% aligned to likely contaminants (14% Primates, 16.5% Birds, 0.5% *Felidae*). No plausible diet-associated organism was present at >1% (**Fig. 3b**). Given that we were able to recover multiple segments and in the absence of another possible host, we concluded that the sturgeon was the likely host of this virus.

Phylogenetic analysis of the three polymerase segments revealed that the sturgeon-associated virus consistently falls within the influenza clade but is basal to all known influenza viruses (**Fig. 3c-e**). Previously, the most divergent virus in the influenza clade was an influenza-like virus associated with a hagfish^12^. However, our analysis demonstrated that this placement is dependent on the alignment method. Alignment with MAFFT recapitulated the reported placement of the hagfish-associated polymerase segments when the sturgeon-associated virus was added to the data set (**Fig. S4**). In all three cases, both hagfish- and sturgeon-associated polymerase segments were basal to known influenza viruses, with the sturgeon segments falling basal to those of the hagfish. In contrast, alignment with MUSCLE rendered the placement of the hagfish-associated segments less certain, placing the PB2 and PA segments basal to influenza A and B viruses (**Fig. 3d,e**). The polymerase segments of the sturgeon-associated virus were basal to known influenza viruses when either alignment method was used. Moreover, the placement of the PB1 and PB3 segments of our sturgeon-associated virus was relatively well supported at the base of the influenza clade (ufboot = 76 and 92, respectively, **Fig. 3c,e**).

The uncertainty around the placement of the hagfish-associated polymerase segments combined with the phylogenetic relationship of the hagfish- and sturgeon-associated segments contradicts strict virus-host codivergence. Hagfish (Agnatha, jawless vertebrates) are phylogenetically basal to sturgeon (Actinopterygii). However, our findings do suggest that influenza viruses can infect all classes of fish such that these animals may have served as early, if not the first, hosts of influenza virus before it spilled over into mammals.

### Phylogenetic relationship of Crustacea- and Chelicerata-infecting *Quaranjaviridae* suggests a history of host jumps

Having demonstrated that aquatic animals were likely central to the evolution of the *Articulavirales*, we next investigated their role in the evolution of the *Quaranjaviridae*. Nearly all viruses in this family, including those identified in this study, are associated with arthropods as their primary host. While we identified a quaranja-like virus in a horsehair worm (Nematomorpha), as this animal parasitizes arthropods, we assumed that the respective virus was more likely to be associated with an arthropod than with the worm. Previous studies have also identified *Quaranjaviridae* in sediment^14^ and faeces samples^35^, and host associations cannot be determined in these instances. Among the 49 novel *Quaranjaviridae* we identified here, 7 were associated with crustaceans (1 shrimp, 2 crab, 1 squat lobster, 1 beach hopper, 1 copepod, 1 tanaid).

To evaluate the relationship of these aquatic viruses within the family, we inferred maximum likelihood phylogenetic trees for the three polymerase segments, the nucleoprotein (NP), and hemagglutinin (HA). The composition of these trees necessarily differed as not all segments (particularly NP and HA) were not publicly available for known viruses preventing direct topological comparisons. PB1 segments were available for all viruses, rendering this tree arguably the most reliable.

The topology of the PB1 phylogeny reveals patterns of both virus-host codivergence and host jumping. This phylogeny can be divided into two clades (**Fig. 4a**). Clade 1 comprises predominantly Hexapoda-associated viruses. However, all four Arthropoda subphyla are represented in this clade, and their relative placement is consistent with the host phylogeny, in which Chelicerata (i.e., ticks, mites, and scorpions) are thought to have evolved first while Crustacea are more closely related to Hexapoda^36^ (**Fig. 4a, *inset***). In this study, we identified a virus associated with centipede (Myriapoda), and this virus falls between Chelicerata- and Crustacea-associated viruses, all of which are basal to Hexapoda-associated viruses (**Fig. 4a**). The two Crustacea-associated viruses in this clade were identified in Malacostraca (tanaid and Pachyschesis). Viruses associated with the same host genera within Hexapoda were generally more closely related suggesting limited recent cross-species transmission. For example, mosquito-associated viruses largely clustered together (**Fig. 4a**).

**Figure 4.**
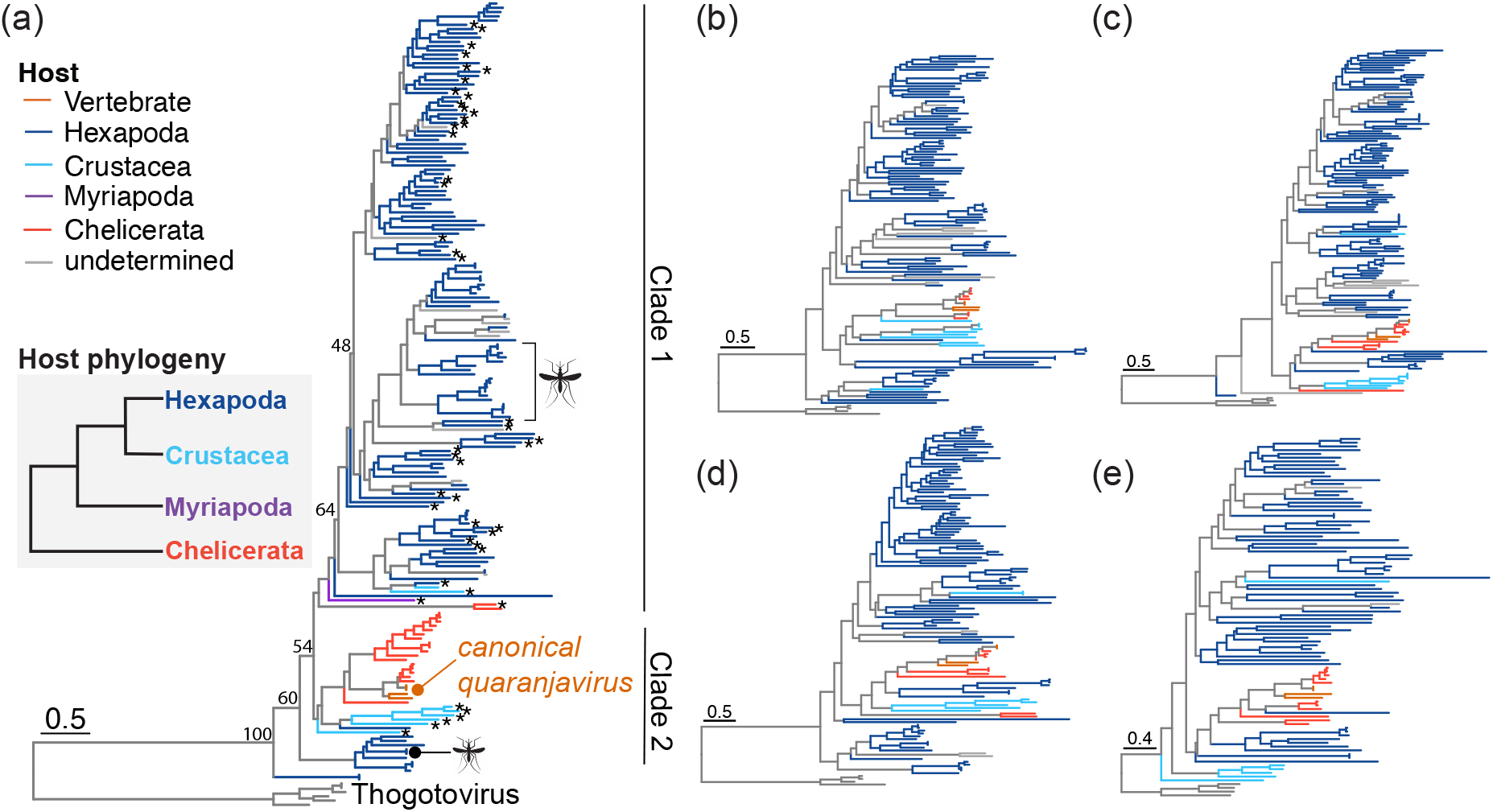
Evolution of the *Quaranjaviridae* is inconsistent with virus-host codivergence. (a-e) Maximum likelihood phylogenies of the *Quaranjaviridae* PB1 rooted with *Thogotovirus*. (a), PB2 (b), PA (c), NP (d), HA (e). Each phylogeny was rooted using thogotovirus as an outgroup. Branch colour indicates putative host. Due to limitations of data availability, differing numbers of segment were used for each tree (PB1: n = 180; PB2: n = 132; PA: n = 146; NP: n = 127; HA: n = 96) (a) Mosquito-borne viruses are denoted with a mosquito icon. Host phylogeny adapted from Thomas *et al*.^36^ *Novel viruses identified in this study. Branch lengths are scaled according to the number of amino acid substitutions per site.

In contrast, the organisation of Clade 2 is consistent with a history of host jumps. Segments from six Crustacea-associated viruses (2 crab-associated, 1 lobster-associated, 1 shrimp-associated, 1 beach hopper-associated, 1 copepod-associated) form a sister clade to Chelicerata-associated quaranjaviruses, and this relationship was observed across multiple segments (**Fig. 4a-d**). The only exception was the HA phylogeny, in which Crustacea-associated viruses form a basal clade to the family (**Fig. 4e**). However, due to the limited availability of HA segments in public data sets, this placement is likely to be an artefact of undersampling. Not only is this topology inconsistent with codivergence, it is suggestive of substantial host plasticity within a short genetic distance. This plasticity is further demonstrated by the canonical quaranjaviruses (Quaranjfil quaranjavirus, Johnston Atoll quaranjavirus) because they are the only known *Quaranjaviridae* to also infect vertebrates (**Fig. 4a, *“canonical quaranjavirus”***). Lastly, it is notable that Chelicerata- and Crustacea-associated viruses share a close phylogenetic relationship as this may reflect an aquatic-terrestrial transition as has been observed throughout the order.

## DISCUSSION

We present evidence that the order *Articulavirales* first emerged in aquatic ecosystems and should be reorganised into four families. We document the first evidence of Cnidaria-associated *Articulavirales*, demonstrating that viruses in this order are likely able to infect basal invertebrates. In addition to the identification of this novel family (the *Cnidenomoviridae*), we greatly expanded the number of known quaranja-like viruses and propose that this genus be reclassified as a family (*Quaranjaviridae*).

Although accurately rooting the *Articulavirales* phylogenetic tree is challenging, basal invertebrates rather than fish are likely to be among the first hosts of the *Articulavirales*. This topology avoids introducing vertebrate-invertebrate transitions. Furthermore, if the *Amnoonviridae* is the root of the tree, the most parsimonious host progression places fish as the first hosts of the *Orthomyxoviridae*. The PB1 segment of TiLV is about 700nt shorter than the PB1 segments within *Orthomyxoviridae*. Additional segments have not been characterised for TiLV, but the PB2 and PA segments of salmon isavirus are approximately 150nt and 300nt shorter, respectively, than their counterparts in the *Orthomyxoviridae*. This relationship implies that *Articulavirales* segments have increased in length over time, contradicting some theories of virus genomic architectural evolution^9^. Moreover, the PB2 segment of the novel sturgeon-associated influenza virus is notably longer than the PB2 segments from descendant taxa in the clade, implying a lengthening and then shortening of this segment. For these reasons, we argue that the *Cnidenomoviridae* is the most parsimonious basal group within the currently available *Articulavirales* phylogeny. Importantly, however, both topologies support an aquatic origin of the *Articulavirales* and indicate that this order may have persisted since the split of Anthozoa and Medusozoa Cnidaria in the Ediacaran (~640 million years ago).

Consideration of the *Articulavirales* as a whole highlights that viruses within this order utilise a large repertoire of transmission routes. Influenza virus spreads via respiratory droplets in mammals and faeces among birds. Fish-fish influenza transmission may also be respiratory, as fish- and amphibian-associated influenza viruses have been detected in gill tissue^23^. There is evidence that both salmon isavirus and TiLV can be transmitted vertically^37,38^ as well as horizontally, potentially through an oral-fecal route^39^. Characterising the transmission dynamics of the *Cnidenomoviridae* between basal invertebrates is beyond the scope of this study, but previous studies have suggested that crustaceans could serve as vectors for aquatic RNA viruses^40^. Invertebrate-vertebrate transmission facilitated by terrestrial arthropods has evolved independently at least twice within the *Articulavirales* (quaranjavirus and thogotovirus), and motile animals would have been necessary if the *Articulavirales* first emerged in sessile invertebrates such as corals. Crustaceans could play such a role, but fish are also plausible vector candidates because they can travel long distances and serve as hosts for at least two *Articulavirales* families. Answering this question and elucidating the mechanism by which the *Quaranjaviridae* spread among non-vertebrate biting arthropods will require experimental data but could shed light on mechanisms of cross-species transmission.

Understanding aquatic transmission dynamics could have implications for virus emergence in terrestrial vertebrates because our findings support the hypothesis that fish were early, if not the first, hosts of influenza virus. While our results do not support a strict virus-host codivergence model due to the relationship of the sturgeon- and hagfish-associated influenza viruses, the presence of influenza-like viruses in both animals suggests that this virus has permeated throughout the entire fish phylogeny. Cyclostomata, to which Myxini (including hagfish) belong, is thought to have diverged from other vertebrate phyla around 600 million years ago^41^, suggesting that *Articulavirales* adapted to vertebrate hosts early in its evolutionary history. No influenza-like viruses have been detected in invertebrates. Although substantial sampling gaps persist, the discovery of other *Articulavirales* in a wide range of invertebrate hosts suggests that influenza, as defined as a distinct phylogenetic group, is limited to vertebrate hosts, and implies that fish were the first. A similar evolutionary history has been proposed for the vertebrate-infecting family *Coronaviridae*^42^. Amphibians form an evolutionary lineage between fish and reptiles and bridge aquatic and terrestrial ecosystems. The relative placement of amphibians is consistent with both virus-host codivergence and an aquatic-terrestrial transition. We find that the latter is more strongly supported by the phylogeny of the influenza clade. Taken together, the evolution of influenza viruses may have been shaped by cross-class host jumps, which are still observable today (e.g., bird-to-mammal transmission of H5N1^43^).

Our study places vector-borne virus emergence into an evolutionary context. In the family *Quaranjaviridae*, only arachnid-associated viruses are known to spill over into vertebrates. While this could be due in part to the biting behaviour of ticks, the *Quaranjaviridae* have also been detected in mosquito genera known to transmit RNA viruses to humans (*Aedes* and *Culex*). Mosquito-borne transmission of quaranjaviruses to humans has not been reported. It may be that cryptic transmission does occur; however, given that vertebrates and particularly humans are disproportionately oversampled, these events are likely to be captured if they are associated with overt disease. Thus, either spill over of mosquito-borne quaranjaviruses occurs but does not cause disease or spill over of these viruses into vertebrates is evolutionarily prohibited. If the former, it may be that these viruses are unable to evade the vertebrate adaptive immune system. If the latter, they may not be able to adapt to vertebrate host receptors. In either case, the absence of mosquito-borne quaranjavirus conferred disease suggests that there are key molecular differences between Hexapoda-borne and Chelicerata-borne quaranjaviruses, and the latter are better able to adapt to vertebrate hosts. Elucidating these differences will require experimental data and could have important implications for understanding vector-borne virus emergence in vertebrates. The close relationship between arachnid- and crustacean-associated Quaranjaviridae again indicates that the evolutionary history of the *Articulavirales* has not followed strict virus-host codivergence. Chelicerata are basal to both Hexapoda and Crustacea^36^, yet Crustacea-associated viruses are more closely related to Chelicerata-associated viruses and are basal to the rest of the family. Sampling horseshoe crabs, which are ancient, aquatic Chelicerata, could provide insight into how this transition occurred.

This study was not without limitations. We utilised publicly available data for most of our analyses, introducing sampling biases (e.g., overrepresentation of Arthropoda hosts) and limiting the size of our data set. As such, the range of hosts of the *Articulavirales* is likely far larger than what we have described. Additionally, the use of cross-sectional metagenomic data precludes the assignment of true virus-host associations. Despite this, the associations we assumed were supported by phylogenetic analysis. For example, viruses identified in *Heliopora* corals cluster together, as do most mosquito-associated *Quaranjaviridae*. Finally, while the overall topology of the phylogenies we inferred had strong bootstrap support, internal tree nodes were less well supported. This is likely due to substantial undersampling within the order, and the addition of more viruses when discovered should improve the veracity of the placement of internal nodes. For this reason, we avoided overly interpreting internal relationships, which limited the depth of our conclusions.

We have demonstrated the power of metagenomic data in elucidating the origins of viruses that are the focus of modern public health initiatives. Continuing to explore virus diversity in a wide range of animal hosts will help to answer key evolutionary questions. For example, did the evolution of the adaptive immune system coincide with incidence of virus-associated overt disease? Answering this and related questions will be especially useful for improving methods to estimate zoonotic risk. Importantly, our findings show that, as the first animals to evolve on Earth, aquatic Metazoa have played a central role in shaping the global virome. Further exploration of marine virus diversity will almost certainly yield deeper insights into the drivers of virus diversity and shed light on how these simple replicating entities have successfully parasitised all known forms of life.

## METHODS

### Sample collection

Both coral samples were collected and identified by Dr. Zoe Richards at the following locations: *Heliopora coerulea* #210, WAM Z42167, subtidal 12m, collected 20.09.2016, NW Montelivet Island, Bonaparte Archipelago, inshore Kimberley, Western Australia. Site 196; S14.28778, E125.2127. *Acropora samoensis*, WAM Z421003, #158, intertidal, collected 18.09.2016, North Patricia Island, Bonaparte Archipelago, inshore Kimberely, Western Australia. Site 189S14.266603, E125.29700.

### RNA extraction and sequencing

The first coral (*Heliopora coerulea*) was processed in 2018. The sample was thawed on ice, removed from RNA later, and rinsed gently with sterile RNA and DNA-free 1X PBS solution (GIBCO). Sterile tools were then used to remove a small fragment (~1g), with care taken to include both soft and hard tissues. The tissue was homogenised using a sterile drill, and total RNA was extracted with the RNeasy Plus Mini Kit (Qiagen). RNA quality was determined using Bioanalyzer fragment analysis, followed by Ribo-Zero Plus library preparation and nextgeneration sequencing (Illumina HiSeq 2500) performed at the Australian Genome Research Facility (AGRF).

The second coral (*Acropora samoensis*) was processed in 2022. This sample was homogenised in liquid nitrogen using a mortar and pestle. RNA was extracted using the RNeasy Plus Mini Kit (Qiagen). The quality and concentration of the RNA were assessed with Qubit. Ribo-Zero Plus library preparation and next-generation sequencing (Illumina NovaSeq S4) were performed at AGRF. Negative extraction controls were used to assess potential contaminants. No relevant contaminants were identified.

### Virus discovery

Raw sequencing reads were trimmed using Trimmomatic v0.38^44^. Reads mapping to rRNA were removed using sortMeRNA v4.3.3^45^. Contigs were assembled from trimmed and filtered reads using MEGAHIT v1.2.9^46^. Assembled reads were screened against the NCBI non-redundant (nr) protein database (as of June 2022) and a custom RdRp database using Diamond BLASTx v.2.0.9. The custom RdRp database contains all currently identified RdRp sequences. In both cases, we used an e-value cut-off of 1×10^−5^. Contigs with hits to known viruses with less than 80% amino acid identity were considered new virus species. All contigs with hits to the RdRp database were cross-checked against the nr results, and virus candidates that hit to host genes were excluded from further analysis. Contigs identified as *Articulavirales* segments were translated using the Expasy online translation tool (https://web.expasy.org/translate/), and each translation was visually inspected in Geneious. Read abundance was estimated using RSEM v.1.3.0^47^ implemented in Trinity v.2.5.1^48^.

### Assessment of endogenous virus elements (EVEs)

All virus candidates were compared to the nr database (as of June 2022) to confirm that they did not share similarity to known host genes. We also assessed the presence of host gene contamination using the contamination function implemented in CheckV^49^. No virus candidates were found to be EVEs.

### NCBI Transcriptome Shotgun Assemblies (TSA) database screen

We performed tBLASTn searches limited to the following organisms: Ascidiacea, Demospongiae, Cnidaria, Arthropoda, Bryozoa, fish (Agnatha, Chondrichthyes, Actinopterygii, Osteichthyes actinopterygian, Osteichthyes coelacanthiform, Osteichthyes dipnoan, Hyperoartia, Gnathostomata, Petromyzontidae, marine lamprey, Arctic lamprey), and Amphibia. These organisms encompass the majority of aquatic animals with available TSA libraries. To identify divergent PB1 segments, we used Wenling hagfish influenza-like virus (AVM87635), Beihai orthomyxo-like virus 2 (APG77864) as input, as well as both PB1 segments identified in corals. Divergent hits (less than 80% amino acid identity) were confirmed to be novel using BLASTx (nr database). Each was translated using Expasy and visually inspected in Geneious. In all cases, only translations containing the RdRp SDD motif that defines the *Articulavirales* were considered. Two contigs were removed due to mutations in this motif. A preliminary phylogenetic analysis was performed to remove identical sequences. In this case, sequences were aligned in MAFFT v7.490^32^, and the phylogenetic tree was inferred using the maximum likelihood approach in IQ-TREE v1.6.12^30^ with ModelFinder, which selected LG+F+R10 as the best fit model.

### Serratus screen

To identify contigs assembled as part of the Serratus project^27^ containing “novel” or unassigned *Orthomyxoviridae* PB1 segments, we downloaded and manually queried the master RdRp contig and metadata file for *Orthomyxoviridae*-like PALMdb palmprint (or barcode) accessions (n=336). These PALMdb palmprints are assigned based on taxonomy and clustered based on 90% identity^50^. Putative novel Orthomyxo-like PB1 sequences were manually validated using BLASTx against the NCBI nr virus database. SRA accessions from identified novel viral-like sequences were assembled using MEGAHIT, and additional segments were extracted from assemblies using tBLASTn.

### Phylogenetic tree estimation

To compile a representative data set of publicly available *Articulavirales* PB1 segments, we downloaded all non-influenza *Articulavirales* PB1 segments available on NCBI Virus that were at least 500 amino acids in length (n = 291). For completeness, we used all available spellings of ‘PB1’ and ‘polymerase basic 1’. Sequences associated with a laboratory host were excluded. We included all influenza-like viruses identified in non-mammalian and non-reptilian hosts (n = 8), and a representative sample of influenza A, B, C, and D viruses (n = 22). Identical sequences were removed using CD-HIT v4.6.1 with a threshold of 0.99. The remaining sequences were aligned with MUSCLE v5.1^29^. After trimming the aligned sequences to the conserved motifs in Geneious and removing gaps with trimAl v1.4^51^, identical sequences were again removed according to genetic distance. The final data set contained 288 sequences.

Sequences for all other phylogenies presented in this study were aligned using MUSCLE v5.1 unless otherwise specified. In all cases, gaps were removed using trimAl v1.4., and maximum likelihood phylogenetic trees based on amino acid sequences were inferred in IQ-TREE v1.6.12 using 1000 ultrafast bootstraps and ModelFinder (**Table 1**). Trees were rendered with ggtree^52^ implemented in R v4.1.2 and visualised with Adobe Illustrator.

**Table 1:**
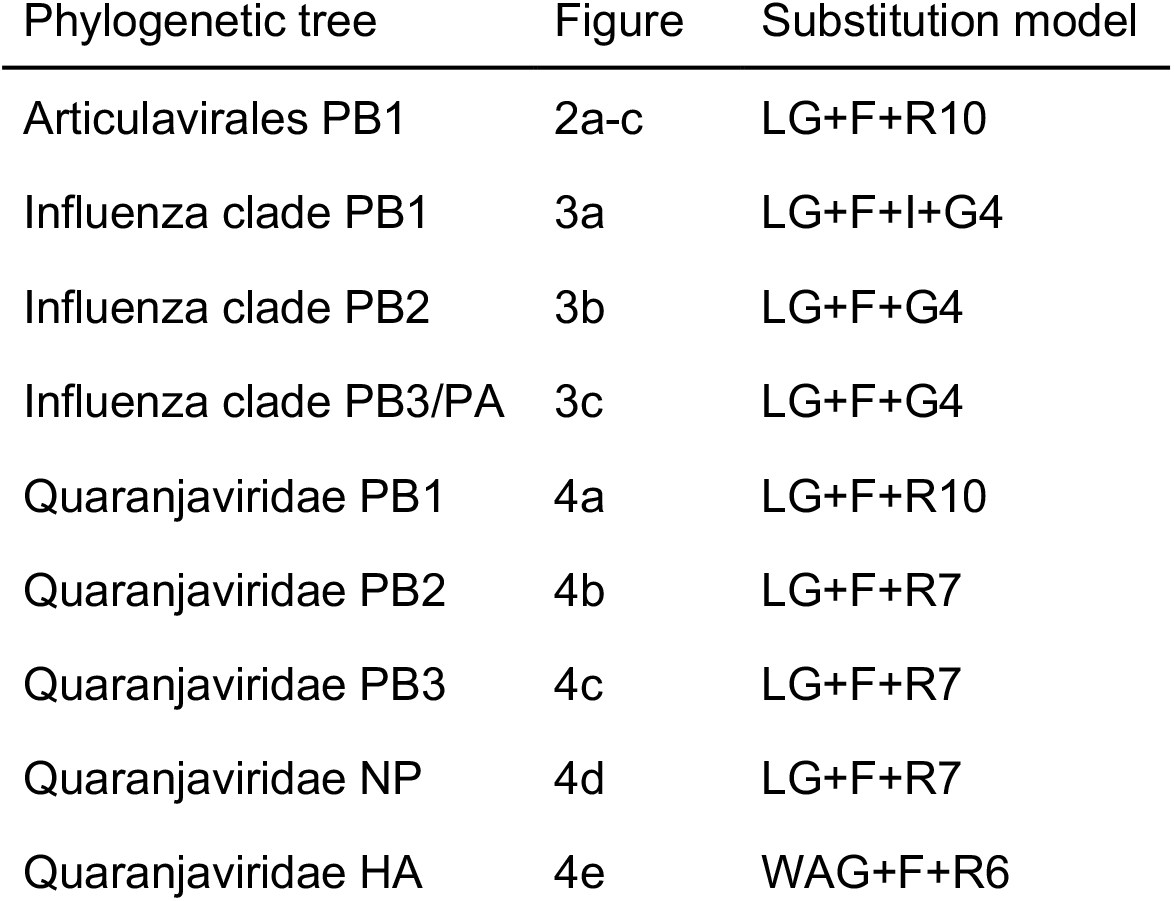
Substitution models selected by Model Finder during phylogenetic inference.

### Analysis of host gene distribution in sequencing libraries

We analysed the composition (defined by host gene assignment and distribution) of the sequencing libraries associated with the Siberian sturgeon (*Acipenser baerii*, TSA: GIPE01, BioProject: PRJNA591120) and the corals from which we extracted RNA (*Acropora samoensis* and *Heliopora coerulea*). For the sturgeon library, the transcriptome shotgun contigs preassembled and associated with the BioProject ID were used as input. In all three cases, we used KMA v1.3.9a^33^ and CCMetagen v1.1.3^34^ and visualised the results with Prism v9.5.0 and Adobe Illustrator.

## ACKNOWLEDGMENTS

We thank Prof. Alexander Sens for his suggestion of the name *Cnidenomoviridae*. The name derives from the Greek Κνίδη δ (nettle, or sea nettle, and the root of Cnidaria) and -νόμος (’dwelling in or ‘pasturing on’). We also thank J-S Eden and C Le Lay for their assistance extracting RNA from the *Heliopora* coral. This work was funded by a National Health and Medical Research Council (NHMRC) Investigator award and by AIR@InnoHK administered by the Innovation and Technology Commission, Hong Kong Special Administrative Region, China. Coral samples were collected on fieldwork funded by the Western Australian Museum, Woodside Energy, the Net Conservation Benefits Fund and ARC Linkage Project LP160101508.

## AUTHOR CONTRIBUTIONS

MEP and ECH conceptualised this study. ZTR collected the coral samples. MEP, RP, JCO, KVB, and IV performed the analyses. MEP wrote the original manuscript draft. MEP, RP, JCO, KVB, IV, ZTR, and ECH edited and revised the manuscript. ECH supervised the project.

## Notes

### Competing Interest Statement

The authors have declared no competing interest.

