## Supplementary material for "Evidence for an aquatic origin of influenza virus and the order *Articulavirales*": Figure S3

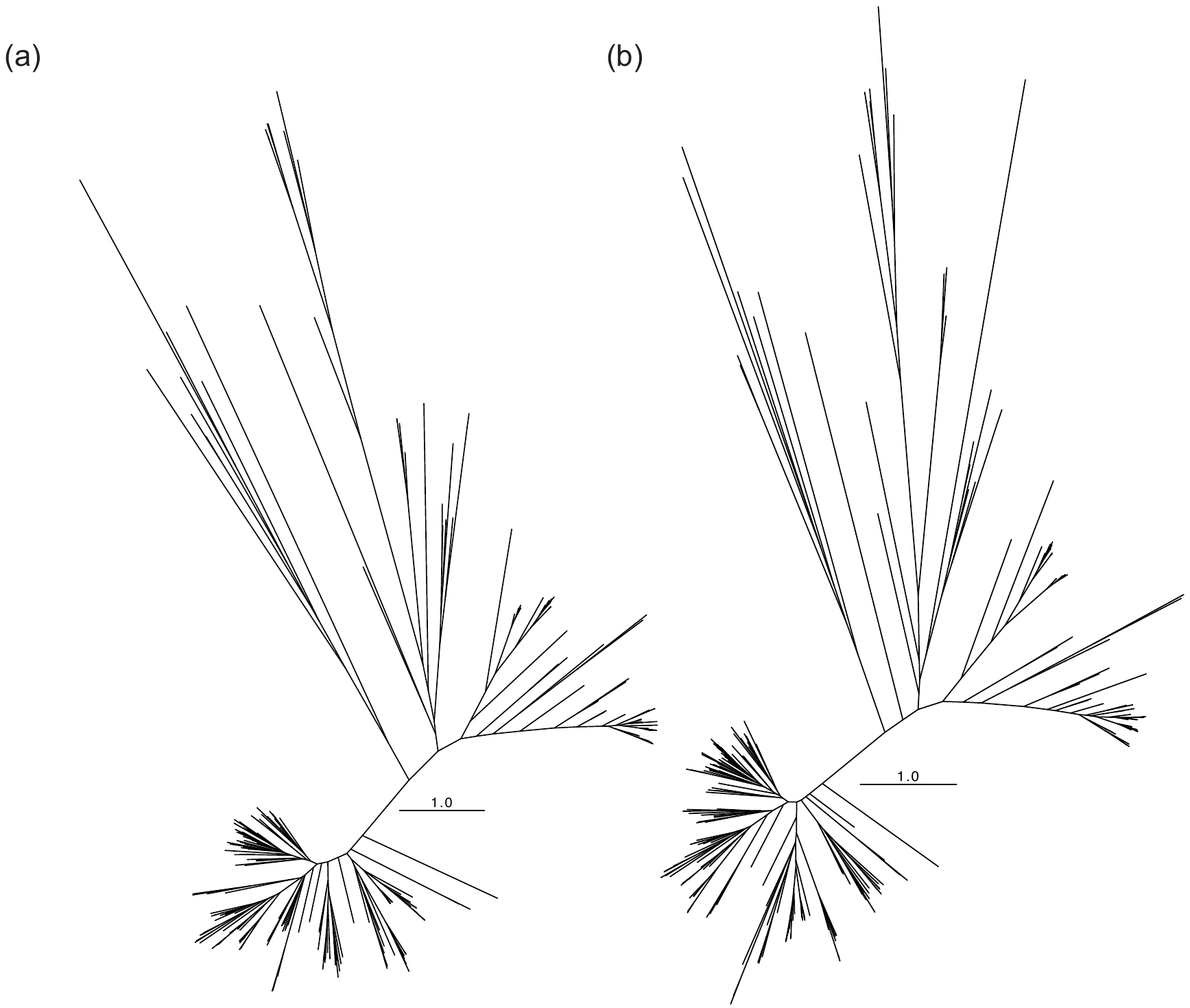

**Figure S3. Comparable tree topologies inferred when different sequence alignment methods were used.** *Articulavirales* tree (**Fig. 1**) aligned with MUSCLE (a) and MAFFT (b).
