## Supplementary material for "Evidence for an aquatic origin of influenza virus and the order *Articulavirales*": Figure S2

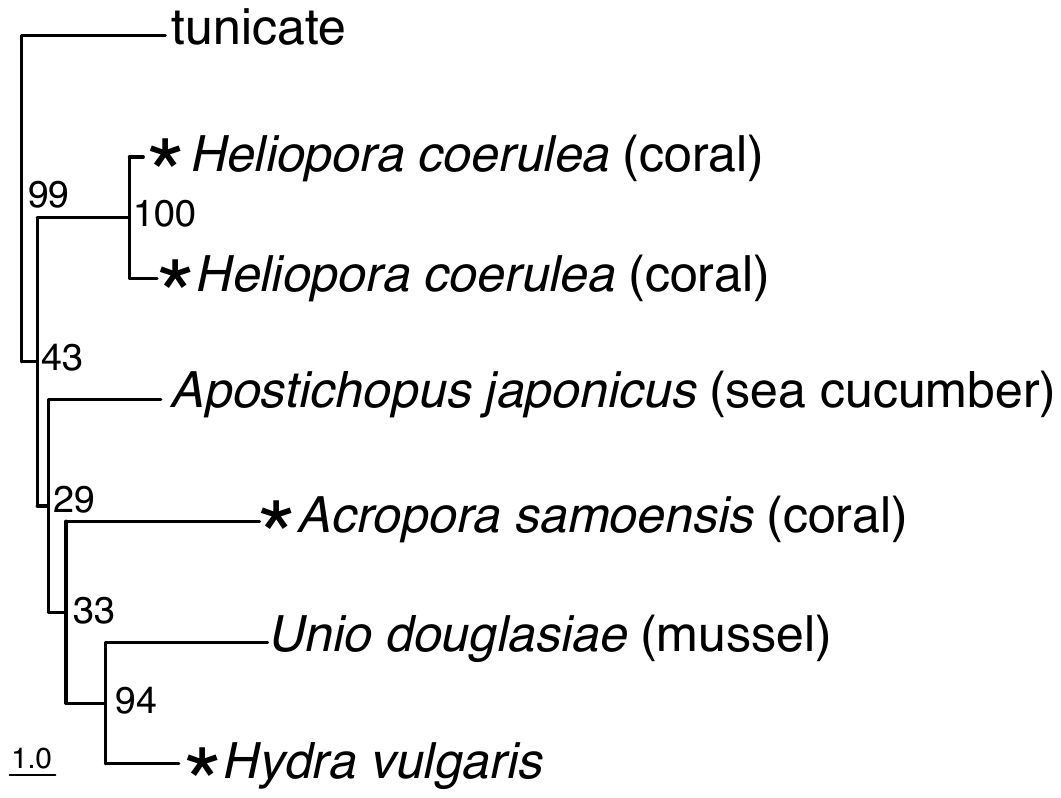

**Figure S2. Cnidenomoviridae clade extracted from rooted Articulavirales phylogenetic tree (Fig. 1).** *Denotes viruses identified in this study
