## Supplementary material for "Evidence for an aquatic origin of influenza virus and the order *Articulavirales*": Figure S1

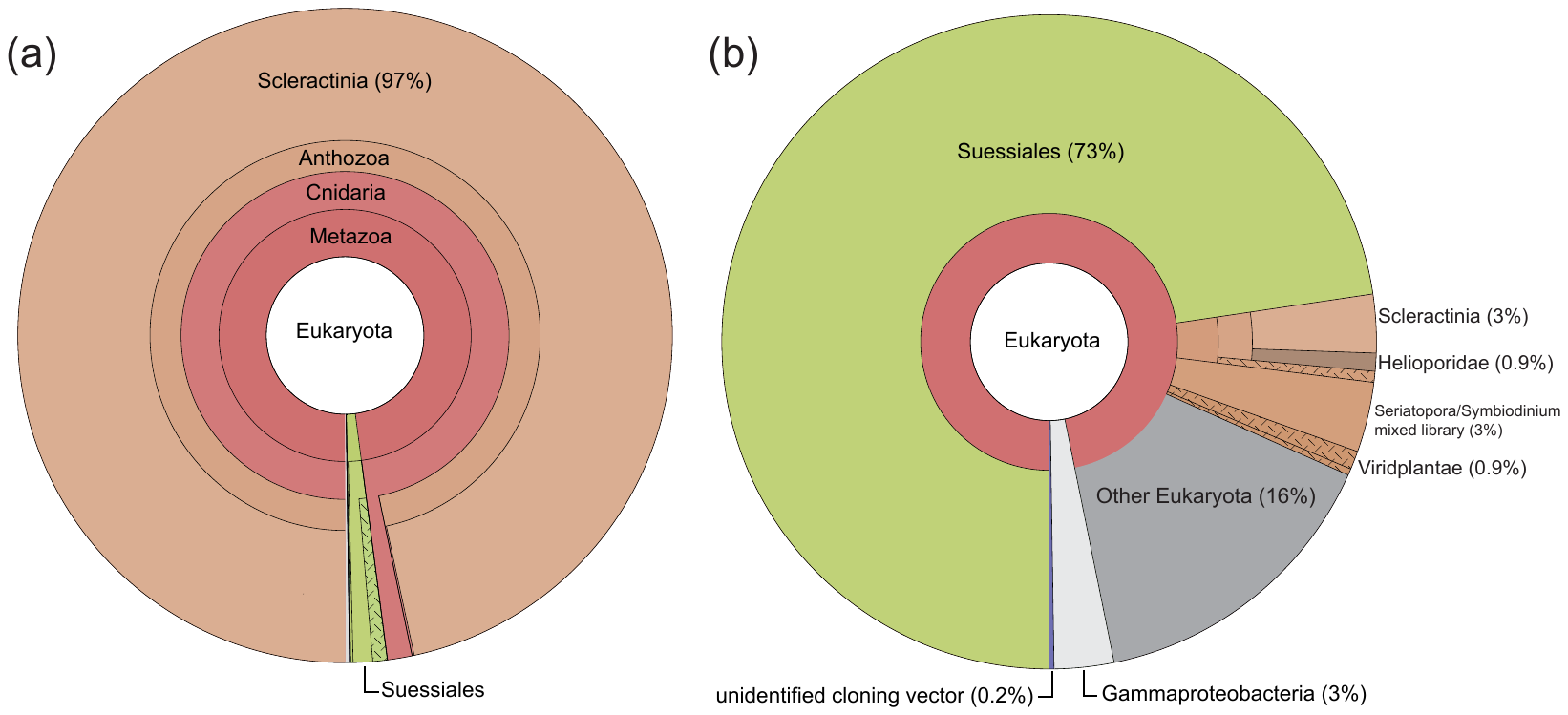

**Figure S1. Composition of coral sequencing libraries derived using KMA and CCMetagen.** (a) *Acropora samoensis* library (b) *Heliopora coerula* library
