## Supplementary material for "Evidence for an aquatic origin of influenza virus and the order *Articulavirales*": Table S1

**Table S1. Summary of BLAST results for novel coral-associated *Articulavirales*.** nr: non-redundant protein database (NCBI), custom: custom RdRp database

| Host species | BLAST (nr/custom) | % identity (nr/custom) | e-value (nr/custom) |
| --- | --- | --- | --- |
| *Heliopora coerulea* | PB1, Beihai orthomyxo-like virus 2/PB1, Ornate chorus frog influenza-like virus | 22.6/23.7 | 7.14e-15/2.66e-14 |
| *Acropora samoensis* | PB1, Soybean thrips quaranja-like virus 2/none | 25.2/none | 9.18e-06 |
